# Comparative attractiveness of *Anopheles quadriannulatus* and *Anopheles arabiensis* to humans estimated by comparing the relative abundance of these two species in larval samples, unbaited adult catches and human-baited adult catches

**DOI:** 10.1101/2024.08.07.606970

**Authors:** Deogratius R Kavishe, Katrina A Walsh, Rogath V Msoffe, Lily M Duggan, Lucia J Tarimo, Fidelma Butler, Nicodem J. Govella, Emmanuel W. Kaindoa, Gerry F Killeen

## Abstract

Understanding mosquito vector behaviour, abundance, and bionomics is crucial for effective malaria prevention. Since most malaria parasites in humans are strict anthroponoses, mosquito preference for humans as a blood source is a key determinant of transmission intensity and intervention strategies. This study compares the attraction of *Anopheles arabiensis* and *Anopheles quadriannulatus* to humans by assessing their relative abundance in larval samples and adult mosquito catches using unbaited and human-baited traps.

The research investigated how the abundance of these sibling species varies with the local availability of humans, livestock, and wildlife as potential blood sources. Surveys of larval and adult mosquito populations were conducted at 40 mobile camping locations distributed across a landscape mosaic of different habitat types, with a gradient of land use practices ranging from comprehensive conversion to agriculture and human settlement to essentially intact natural ecosystems inside well-protected conservation areas. Larvae were collected from all accessible water bodies within a 2 km radius of each mobile camp, while adults were surveyed using four light traps and one interception netting barrier trap at each camp. Light traps were strategically placed at locations such as beside a human-occupied tent, near the camp, in a nearby streambed, and in an open natural glade, while the netting barrier trap was placed in the open natural glade.

Most adult *Anopheles gambiae* complex mosquitoes caught were unfed and presumably host-seeking. The mean catches of the complex in light traps next to the human-occupied tent was over four times higher than elsewhere around the camp (p<<0.0001). Catches decreased with distance from humans, suggesting attraction to humans by at least one species in the complex. *An. arabiensis* was caught in greater numbers in the human-occupied tent, while *An. quadriannulatus* catches remained consistently low across all traps, even in wild areas where it was dominant in larval population. The proportion of *An. arabiensis* in adult collections was higher than in larval samples (98.7% versus. 78.3%, p<<0.0001), and adults caught beside human-occupied tents had 36 times higher odds of being *An. arabiensis* rather than *An. quadriannulatus*. Similarly, the barrier trap away from humans but frequently visited by researchers showed a 24-fold enrichment of *An. arabiensis*.

These results confirm the strong attraction of *An. arabiensis* to humans, contrasting with the non-vector *An. quadriannulatus*, which is largely unresponsive to humans. Light traps by human-occupied tents efficiently capture anthropophagic mosquitoes outdoors, while unbaited traps far from people appear to give unbiased representations of larval population composition but with very low efficiency. Moreover, netting barriers with human activity attract anthropophagic mosquitoes, turning them into semi-baited traps with moderate efficiency.

## INTRODUCTION

*An. arabiensis* has been described as a stereotypical vector of residual malaria transmission (Killeen et al., 2016) because it exhibits all the plastic behavioural resilience traits that underpin persisting malaria endemicity despite high coverage with long-lasting insecticidal nets (LLINs) and/or indoor residual spraying (IRS) of insecticides (Durnez & Coosemans, 2013; Killeen, 2014; Killeen et al., 2017; WHO, 2014). While it readily attacks people and can thrive while relying largely upon human blood, thus making it an efficient vector of anthroponotic malaria parasites, it can just as readily feed upon cattle, thus allowing it to evade contact with these two key human-centred vector control measures (Charlwood et al., 1995; Gillies & Coetzee, 1987; Killeen et al., 2017; Killeen et al., 2001; Mahande et al., 2007; Meza et al., 2019; Mlacha et al., 2020; Tedrow et al., 2019; Tirados et al., 2006; White, 1974; White et al., 1972). Furthermore, *An. arabiensis* specifically (Killeen et al., 2016), and zoophagic vectors of residual transmission more generally (Durnez & Coosemans, 2013; Killeen, 2014; Killeen et al., 2017; WHO, 2014), also exhibit several other evasive behaviours that minimize their exposure to the active ingredients of LLINs and IRS: outdoor biting, particularly at dawn and dusk when people are often active outdoors, outdoor resting and rapid exit from houses that they enter in search of blood.

*An. arabiensis* may also be described as being neither anthropophagic nor zoophagic, but rather both, because it readily exploits blood from both human and non-human sources (Charlwood et al., 1995; Gillies & Coetzee, 1987; Killeen et al., 2017; Killeen et al., 2001; Meza et al., 2019; Mlacha et al., 2020; Tedrow et al., 2019; Tirados et al., 2011; White, 1974; White et al., 1972). Nevertheless, like most blood-feeding arthropods (Lyimo & Ferguson, 2009), it would be inaccurate to describe this species as a generalist in the strict sense: Thus far, cattle remain the only non-human mammal that they have been found to display a strong, clearly measurable innate affinity for (Costantini et al., 1998; Killeen et al., 2001; Meza et al., 2019; Mlacha et al., 2020; Nguyen et al., 2017; Tedrow et al., 2019), a trait that appears linked to alleles within chromosome inversion 3Ra (Main et al., 2016). Indeed, a recent study in southern Tanzania, to examine the relationship between the distribution of *An. arabiensis* larvae and fine-scale variations in the abundance of people, livestock and wildlife, also identified humans and cattle as the only two mammalian species that were positively associated with increased occupancy of water bodies by this malaria vector species (Walsh, 2023; Walsh et al., In preparation).

Nevertheless, *An. arabiensis* has been presumed to feed upon wild mammals based on their presence within conserved wilderness areas of southern Africa (Braack et al., 1994; Munhenga et al., 2011; Munhenga et al., 2014; Prior & Torr, 2002) even though direct evidence in the form of unambiguously identified bloodmeals are lacking. More recently, apparently self-sustaining populations of this key vector of residual malaria transmission have been documented in the apparent complete absence of humans or cattle, at ecologically intact locations deep inside Nyerere National Park (NNP) in southern Tanzania (Walsh, 2023; Walsh et al., In preparation) However, although larval surveys revealed that *An. arabiensis* was ubiquitous all across this survey area, it was usually outcompeted by *An. quadriannulatus*, another sibling species from within the same *An. gambiae* complex, in the most remote and undisturbed natural ecosystems(Walsh, 2023; Walsh et al., In preparation). In stark contrast to *An. arabiensis*, *An. quadriannulatus* is generally considered to be a non-vector of malaria and far more strictly zoophagic (Gillies & Coetzee, 1987; White, 1974). Indeed, this study in southern Tanzania implicated both impala and bushpig as the only potential blood sources that tipped the competitive balance between these two species in its favour (Walsh, 2023; Walsh et al., In preparation).

As described herein, however, the same study also surveyed adult mosquito populations (Kavishe et al., 2024), yielding samples that were almost exclusively composed of *An. arabiensis*. These adult mosquito data, combined with those from larval surveys at the same time and place (Walsh, 2023; Walsh et al., In preparation), were therefore analysed, to determine whether this key vector species may have been selectively attracted to the human team of investigators, whose camp and the relevant traps were placed in and around.

## METHODS

### Study setting

The ecologically diverse study area illustrated in Figure 1 was ideally suited to addressing the objectives of several complementary conservation biology (Duggan, 2023; Duggan et al., In preparation), vector ecology (Walsh, 2023; Walsh et al., In preparation), and adult mosquito surveys (Kavishe et al., 2024). This integrated set of studies collectively aimed to explore whether *portfolio effects* (Killeen & Reed, 2018) arising from landscape heterogeneities in the availabilities of various potential mammalian blood sources might allow refugia population of malaria vectors to survive in wilderness areas, where anthropogenic selection pressure for insecticide resistance may be greatly attenuated. The pooled adult (Kavishe et al., 2024) and larval (Walsh, 2023; Walsh et al., In preparation) mosquito data are ideal for the secondary analysis presented herein, because the study area for these primary studies (Figure 1) represents a geographical gradient of natural ecosystem integrity dominated by different *An. gambiae* complex sibling species, ranging from fully domesticated land cover with only *An. arabiensis* in the west, through to completely intact natural ecosystems dominated by *An. quadriannulatus* to the east.

**Figure 1.**
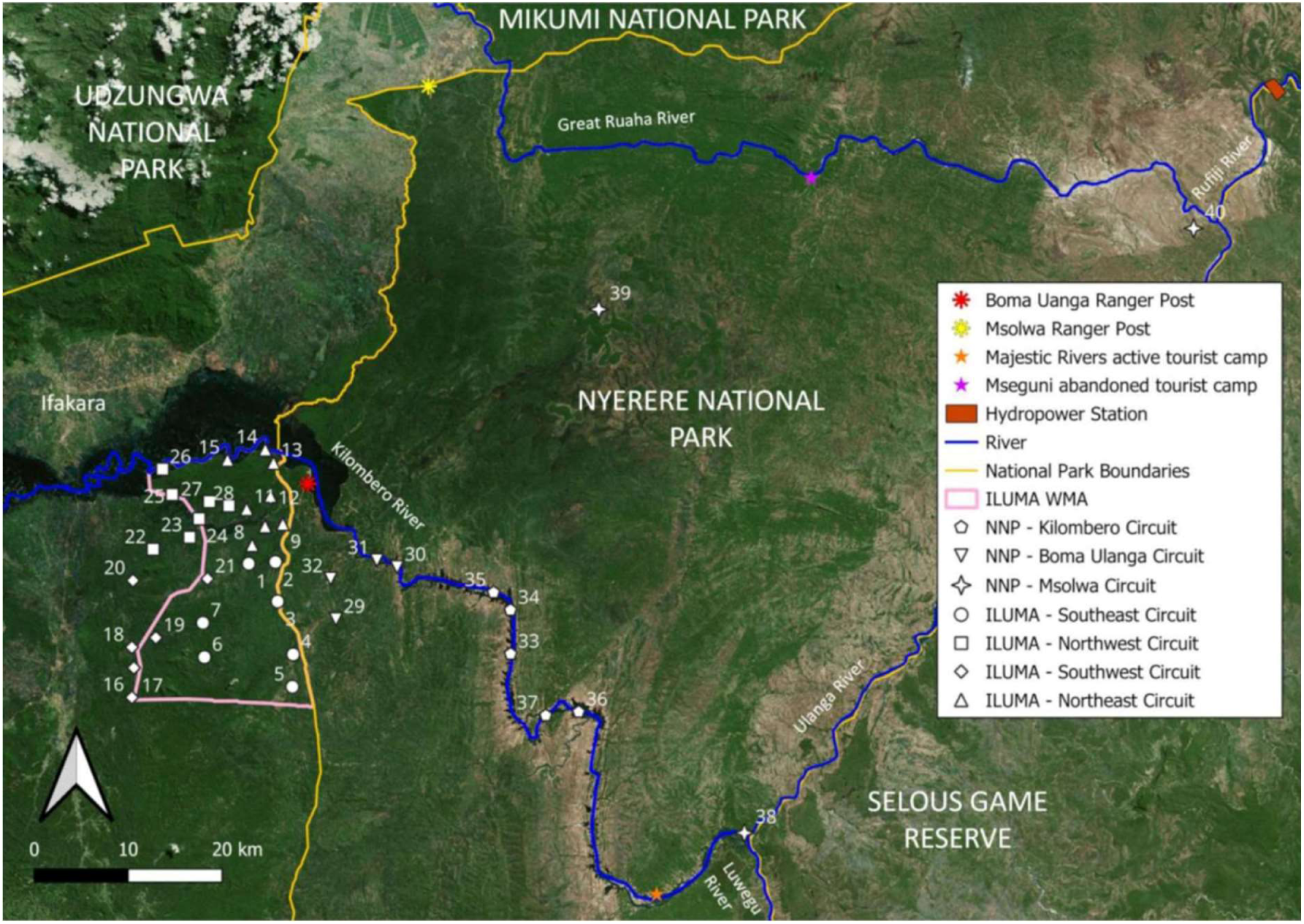
Map displaying the distribution of suitable camping locations used as the sampling frame for all the surveys described in this report (Walsh et al., In preparation). Each of the 40 camps detailed in supplementary file 1 (Walsh et al., In preparation) are illustrated in the geographic context of the boundaries of the Ifakara-Lupiro-Mang’ula Wildlife Management Area (ILUMA WMA), Nyerere National Park (NNP) and the Udzungwa Mountains National Park.

The overall project was carried out using a rolling cross-sectional design, with four rounds of surveys encompassing a total of 40 defined camp locations across NNP, ILUMA WMA, and the villages immediately to the west of it over the course of two years (Figure 1 and supplementary file 1). For the original protocol, a total of 28 camps were selected after scouting potential locations in ILUMA WMA and considering the suggestions of the village game scouts based on their vast personal knowledge and experience of the area. The broad geographic distribution of the camps was planned to encompass as wide range of ecosystem states as possible (Duggan et al., In preparation), by including all parts of ILUMA WMA and the neighbouring domesticated land to the west. However, the exact position of a camp location was ultimately determined by the requirement for perennial surface water from which *Anopheles* larvae and adults could be collected. The presence of one or more glades or valleys with numerous perennial waterbodies, like waterholes, ponds and streambeds near the camp was also required.

Once the camp site was selected, each camp was given a unique number and a name and the full list of camps was then divided into quarterly *circuits* that roughly corresponded to the *southeast*, *southwest*, *northeast*, and *northwest* of the ILUMA WMA, with each circuit named accordingly (Figure 1, supplementary file 1). The circuits were designed to manage the physical challenges of the study, and so subsets of 6 to 8 camps were grouped into distinct circuits based on their geographic proximity to one another and the practicality of visiting them all in a circular route that started and ended centrally at the Msakamba camp. Each circuit had a planned route that minimised walking distances between camps, to ensure that the team had enough time to rest in the afternoon before proceeding with data collection later into the evenings and in the early mornings. Camps were visited consecutively for two nights each and incorporated intensive daily data collection procedures. Therefore, the circuit design for sequentially visiting camps in a rolling cross-sectional survey was crucial for sustaining optimal long-term data collection by limiting investigator fatigue and assuring breaks of a few days in between periods of continuous field work that typically lasted about two weeks for a single circuit.

The original protocol planned for a total of three rounds of sampling to be completed from January to March, March to May, and August to October 2022, representing the short rainy season, long rainy season, and dry season, respectively. It was planned that each round of surveys would incorporate the first 28 camps detailed in supplementary file 1 (Walsh et al., In preparation), but a few camps were omitted from each round for pragmatic reasons, such as a lack of surface water during the dry season or inaccessibility due to severe flooding during intense rains.

However, no camp in the ILUMA WMA appeared to lack signs of human disturbance (Duggan et al., In preparation), and only a few remained relatively well conserved, so four new mobile camps inside NNP were added (number 29-32, figure 1 and supplementary file 1) to the study design at the end of round three in November 2022, forming a new circuit that was surveyed with vehicle support for logistical and safety reasons. These camps were located inside the boundary of NNP, immediately to the east of ILUMA WMA to capture absolute well conserved environments and were accessed by vehicle via the NNP ranger post at *Boma Ulanga*, for which that circuit was named. These were then repeated at the start of the fourth and final round of data collection that was completed from February to July 2023, representing the whole wet season and the beginning of the dry season for that calendar year. Considering the initial unexpected sibling species composition and insecticide phenotype results obtained from the first four NNP camps, it was decided to extend the sampling frame deeper into the park and adjust the field protocol to collect and immediately preserve additional collections of larvae that would not be used to rear adults in the Msakamba insectary (Walsh, 2023; Walsh et al., In preparation)

To obtain samples as far away from human beings, and as deep into well conserved ecosystems as possible, another, eight camps inside NNP were added to the end of the fourth and final round. First, an additional subset of five camps that were accessed by boat via the Kilombero river, and correspondingly named the *Kilombero circuit,* were visited (Figure 1). Pushing even further east into the park, camps 38 to 40 were accessed by vehicle along main park roads via the Msolwa ranger post and were therefore named accordingly as the *Msolwa circuit* (Figure 1, supplementary file 1).

### Adult mosquito surveys

Live adult mosquitoes were trapped using two methods: a netting barrier interception screen, similar to that developed for sampling exophilic mosquitoes in the Pacific (Burkot et al., 2013; Davidson et al., 2018; Keven et al., 2019; Tedrow et al., 2019) and Centres of Disease and Control (CDC) light traps (John W. Hock Company, product number 512). After arriving at a new camp location, a shaded area next to a water source was identified as a suitable camp location for setting up the tents, cooking, resting and processing mosquitoes. The batteries for four CDC light traps that were used for trapping live mosquitoes were then charged as soon as possible using portable solar panels. Also, on overcast days, usually during the rainy season, charged batteries were delivered from the Msakamba central camp (Location 1 in figure 1) once every two days, by the team responsible for picking up the collected mosquito samples and returning them alive to the field insectary (Kavishe et al., 2024). A site within a nearby open valley or natural glade, at an approximate distance of 100 to 200m from the camp, was identified and a five-panel 25m long interception screen made from standard mosquito netting was set up there (Kavishe et al., 2024). Collections at the interception screen were carried out five times after dusk, for every hour between 19:30 and 23:30, and once just before dawn at 05:30, specifically to target the known peak outdoor biting activity of adult *An. arabiensis* in the Kilombero valley (Maia et al., 2016; Russell et al., 2011). Both sides of the screen were scanned in succession, using a torch to carefully inspect each panel in an up-and-down motion for resting mosquitoes. Once seen, each mosquito was suctioned into a collection cup using a custom-made, standard *Prokopack* aspirator powered with a 6V GWESPECS battery (Maia et al., 2011). Separate collection cups were used for every hour and were immediately labelled with the date and time of sampling. A small ball of cotton wool was soaked into a 10% w/v glucose solution and placed on top of the collection cup as a food source.

Additionally, four CDC light traps ran overnight between 19:00 and 07:00, each in a different location to capture any potential variability in host-seeking behaviour or dispersal patterns. Each light trap was set up in one of four distinct ways: One suspended from a tree in an open valley or natural glade far away from humans about 100 meters form the camp, one next to a stream bed 30 to 50 meters from the camp, one within the camp site, and one right next to a human-occupied tent. Traps inside the camp and next to the occupied tent were placed as far away from the campfire as possible. The following morning at 07:00, all captured mosquitoes from the interception screen and the CDC light traps were transferred into clean, labelled paper cups using a mouth aspirator and supplied with fresh glucose solution soaked into cotton wool.

Then an experimental design form template (ED1) from a standardised mosquito collection informatics platform (Kiware et al., 2016) was completed to include variables such as *date*, *camp number*, *collection method*, *habitat type* (referring to where the light trap or barrier screen was placed)*, start time*, *finish time*, *experiment day*, *volunteer initials* and any further *comments* (Supplementary file 2). Samples in the paper cups were labelled with the date and time, as well as the ED1 form *serial number*, *form row* and habitat type values, for sample processing and tracing. These cups containing mosquitoes were then placed into a specifically-designed mosquito carrier backpack and transported alive back to the field insectary at Msakamaba (Kavishe et al., 2024) for further processing.

After the mosquitoes were collected and prepared on the first morning, this procedure was repeated at the same camp on the following evening and morning, after recharging the batteries in the afternoon. On this second morning, the field team would break camp and hike with all the equipment to the next camp location to be surveyed, but only after two Village Game Scouts arriving from the Msakamba central camp delivered supplies and collected the backpack of mosquito samples to return them to the field insectary for sorting, counting, and rearing. After arriving at the new mobile camp location, the routine described above was repeated until the relevant survey circuit was completed.

### Larval surveys

Surveys of mosquito larvae within water bodies at and around each camp were conducted as describe by (Walsh, 2023; Walsh et al., In preparation). The upper limit of 2km from the camp location was selected as the maximum distance at which to survey larvae at, to prevent geographic overlap between habitats surveyed from neighbouring camp locations and to minimise potential spatial autocorrelation effects. Surveys were intensive and conducted in a challenging climate, where habitats were often in open, unshaded areas, so a time limit of four hours was decided upon to mitigate against investigator fatigue and ensure optimal data collection for the full duration of an entire survey circuit of up to 8 camps visited over up to 16 days of continuous field work. This upper limit of survey duration was based on a prior experience of the overall project and an initial pilot of the field procedures, both of which indicated that longer surveys would prove unsustainable, with potential deleterious consequences for both data quality and the surveyor’s wellbeing, especially given the extreme heat in the afternoon. Starting at the camp, surveys were initiated at the nearest waterbodies known to the team of Village Game Scouts who escorted the investigators (Duggan, 2023; Duggan et al., In preparation; Walsh et al., In preparation) and any other potential aquatic habitat that were seen along the way were also surveyed. Although this purposive sampling was obviously not randomised in any way, it proved practically feasible and maximised the number of habitats sampled around each camp in an efficient manner over the limited time frame of four hours. If the 2km distance from the camp was reached before the time limit was up, a different route back to the camp was taken and any potential larval habitats identified along the way were also surveyed.

All larval survey data were recorded using the data entry form provided in supplementary file 3, with the habitat type entered as a numeric code from 1 to 12 that corresponded to twelve broad categories of water bodies. Further information, including a more detailed description of the habitat and the presence or absence of *short vegetation*, *tall vegetation* and *floating vegetation* were recorded by ticking the appropriate column in the relevant row for each habitat. Direct approximate estimates for *water depth* and *perimeter* were made, so that each habitat could be assigned to their corresponding categories on the form (Walsh et al., In preparation).

At each aquatic habitat, small samples of water, each informally referred to in the field as a *dip*, were taken by briefly submerging a standard white 350ml dipper just below the water surface into the habitat at 45-degree angle, so that the suction created by water displacement drew both water and mosquito larvae into the dipper cup.

All dips were taken along the waterbody perimeter and were methodically numbered and spaced according to estimates of the habitat perimeter and criteria defined as follows. If a potential habitat was <2m in perimeter, one or two dips were taken depending on size and feasibility. For human, livestock and wild animal prints, or any other habitat that was too small to be effectively dipped, a standard 280 ml turkey baster was used to suction water and larvae from the habitat into the dipping cup. These habitats were classified as having been dipped once. For medium-sized waterbodies, with perimeters between 2m and 20m, as well as two larger waterbody perimeter categories greater than 20m, and 200m, respectively, the spacing between each dip was estimated by the number of paces taken by the individual carrying out the survey to ensure dips were taken at regular intervals around the full perimeter. Aquatic habitats that were categorised as being approximately 2m to 20m in perimeter, were dipped for a maximum of five times. Therefore, if the habitat was about 10m in perimeter or less, dips were taken once every two paces walked around the perimeter. If the habitat was estimated to be between 10m and 20m, dips were taken once every four paces. For larger waterbodies between 20m and 200m, and those that were greater than 200m, a maximum of 10 dips were taken once every 20 paces and every 50 paces, respectively. These criteria were developed to maximise sampling efficiency, consistency, and productivity after a pilot survey. Even if *An. gambiae* complex larvae were identified immediately, dips were continually taken around the full perimeter or until the maximum number of dips was reached, because as many of these larvae as possible were collected, kept alive, and sent back to the Msakamba field insectary, to be reared for insecticide susceptibility testing.

All field-identified *An. gambiae* complex larvae collected over the first three rounds of surveys, from January to November 2022 were reared to adults for insecticide susceptibility testing (Kavishe et al., Unpublished). Following the observation that the *An. gambiae* population at many camps consisted of a mixture of *An. arabiensis* and *An. quadriannulatus* (Walsh, 2023; Walsh et al., In preparation) as identified by polymerase chain reaction (PCR) (Scott et al., 1993; Wilkins et al., 2006), it was therefore decided to add a supplementary procedure for collecting additional larval samples over the course of the fourth round of surveys, and to preserve these *in situ* to more reliably address these questions about how sibling species composition of the *An. gambiae* complex within larval populations varied with location and potential host availability. The purpose of standardising and limiting the number of dips in the primary larval collection protocol was to strike a compromise between the amount of time spent at each waterbody and the number of different waterbodies that could be surveyed for assessing how occupancy varied across the ecological gradient of the study site. As it was essential to maintain the existing survey protocol without changing it, so that data collection to address the original questions about occupancy and insecticide resistance phenotypes were consistent and comparable, it was decided to supplement those collections of live larvae with more purposive and intensive collections from the same aquatic habitats to obtain larger, more carefully disaggregated samples and preserve them *in situ* for reliable PCR analysis (Scott et al., 1993; Wilkins et al., 2006), yielding data that could be interpreted with far fewer caveats than those derived from larvae reared through to adults (Walsh, 2023; Walsh et al., In preparation).

When a habitat was positively identified for the presence of *An. gambiae* complex, the first individual conducting the occupancy survey continued to collected larvae as per the original protocol that involved the live transportation to the field insectary (Kavishe et al., 2024). However, during this fourth and final round, separate samples of larvae were collected in parallel by a second trained Village Game Scout following immediately behind the first surveyor. This second surveyor collected larvae from aquatic habitats that were positively identified during the survey using the same dipping technique but did so far more exhaustively. Only larvae identified *in situ* as members of the *An. gambiae* complex were immediately preserved by transferring them into a 50ml Falcon^®^ tube filled with 95% ethanol with a clean, disposable 5ml plastic pipette. Directly preserving larvae, right at the aquatic habitat they were collected from, eliminated any potential bias or human error that could distort the sibling species composition during the transport and rearing processes, thus ensuring more robust formal species identification by PCR (Scott et al., 1993; Wilkins et al., 2006) and data that could be more directly interpreted with fewer caveats. This approach also increased the sample sizes to be used for subsequent statistical analyses, as the only focus of the second collector was to identify and preserve as many *An. gambiae* complex larvae as possible from each aquatic habitat found to contain them. Also, given the specific interest in sibling species composition, and likelihood of strong covariance of individual dip samples with habitats, combined with substantial variance among habitats, it was decided to keep separate, specifically labelled samples for each individual habitat, so that anticipated within-habitat covariance and between-habitat variance could be accounted for in the statistical analysis described below with nested random effects.

Unlike the previous surveys, over rounds one to three in 2022, wherein field-identified *An. gambiae* complex larvae from multiple habitats were often pooled together, the larvae from different habitats were all stored as separate batch samples, so that each tube contained preserved larvae from a single aquatic habitat. A target of ten batch samples per camp, taken from ten different larval habitats, was set to enable robust quantification and statistical evaluation of population composition heterogeneity around each camp location. Each habitat-specific batch sample was labelled with the camp number, the serial number and form-row number on the larval surveillance form and given a *breeding site identification number (BSID)* from 1 to 10, so that each batch sample could be traced to a particular aquatic habitat from a particular camp. These batch samples of preserved field-identified *An. gambiae* complex larvae identified based on a characteristic white collar immediately behind the head (Holstein, 1954; Walsh, 2023) were returned to Msakamba and then onward to the Ifakara Health Institute molecular laboratory in Ifakara for PCR analysis (Scott et al., 1993; Wilkins et al., 2006).

### Statistical analysis

All statistical analyses were performed by using the *R*^®^ version 4.1.3 open-source software package, through the *Rstudio*^®^, version 2023.09.1.494 environment (Team, 2022). To estimate mean catches of adult female *Anopheles gambiae s.l*. obtained from light traps placed at four distinct locations around the camp and one interception barrier trap placed far away from it, a generalized linear mixed model (GLMM) lacking an intercept was fitted with the *glmmTMB* package, using a type II negative binomial distribution for the number of mosquitoes from this complex caught as the response variable, while trap type and location was included as the sole categorical predictor variable. A random effect incorporating first-order temporal autocorrelation, using the number of weeks since the study started as the incremental unit of time, nested within the camp location, was used to account for seasonal time trends and seasonally consistent geographical variations across the study area. A similar GLMM with an intercept was fitted to contrast between the light trap beside the human-occupied tent and the rest of the light traps placed farther away from people and from the camp, as well as the interception barrier trap placed further away from it.

Furthermore, to estimate the proportional sibling species composition of the adult *An. gambiae* complex mosquito samples obtained from four different light trap positions, barrier trap, and the larval samples, a logistic GLMM was fitted to the binary sibling species identity outcomes (*An. arabiensis* versus *An. quadriannulatus*) from all PCR amplified specimens from adult and larval collections using the *glmer* function of the *lme4* package. The proportion of the amplified specimens that were *An. arabiensis* was treated as the binomial dependent variable and the collection method as the categorical independent variable. Camp number and date of collection were included as random effects. Similarly, the same GLMM model but with an intercept was fitted to obtain the odd ratio for the proportion of *An. arabiensis* in each of the different adult catches compared to the larval collections.

## RESULTS

A total of adult 22,525 mosquitoes were collected from 797 trap nights of sampling carried out over 240 nights with four light traps and one interception barrier trap. Of these 23.8% (5366) were morphologically identified as *An. gambiae* complex, 1.5% (333) as *An. funestus* group, 13.3% (2995) as other *Anopheles*, and 61.4% (13822) as sundry *Culicines*. The overwhelming majority of female *An. gambiae s.l.* mosquitoes caught (about 96%) were unfed, indicating all these capture methods primarily collected host-seeking mosquitoes (Table 1).

**Table 1.**
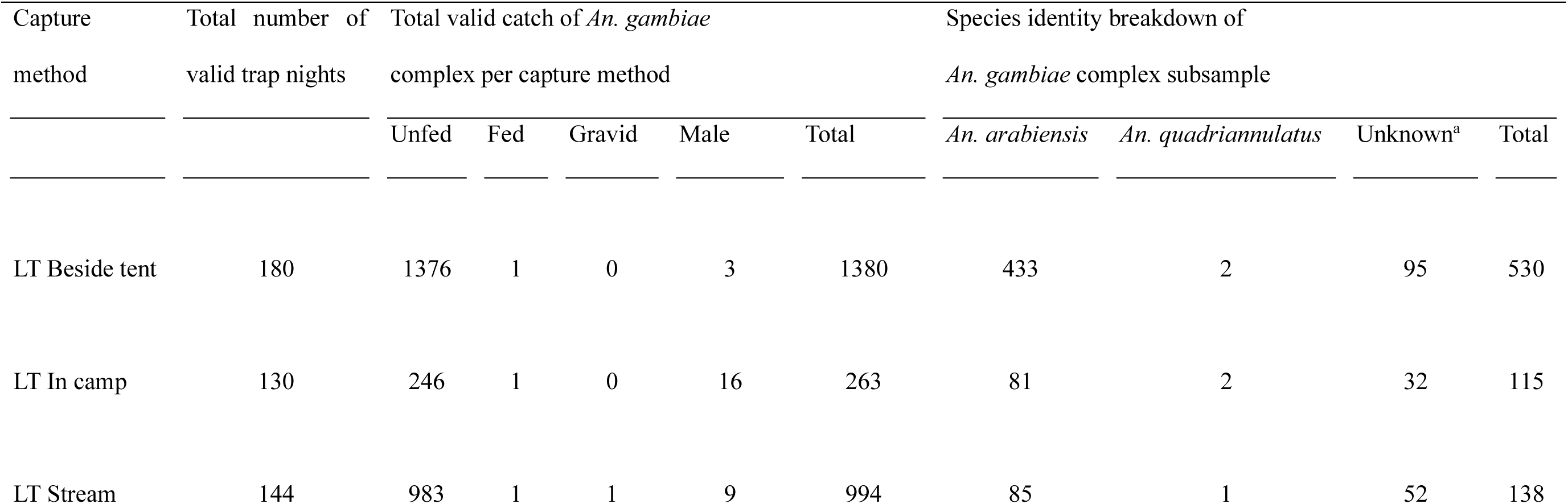
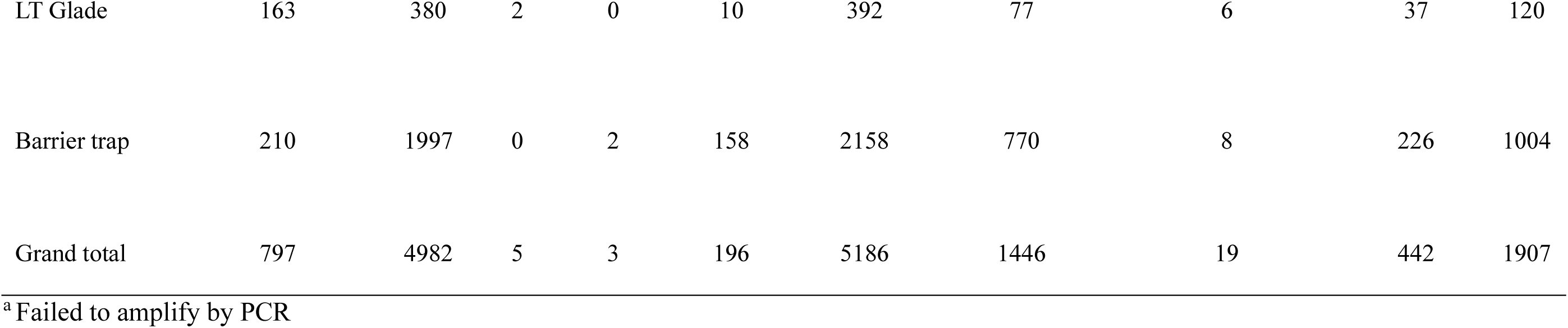
Number and physiological status of adult *An. gambiae* complex mosquitoes captured in 797 valid trap nights in either light traps (LT) placed in one of four distinct locations in and around the camp or one barrier trap. The four different positions used for LT were as follows (1) beside a tent occupied by one or two humans, (2) close to the camp, (3) in a nearby streambed, (4) in an open natural glade. The one netting barrier interception trap placed in an open natural glade same as the fourth light trap. Also, PCR results for subsamples of *An. gambie s.l.* are presented broken down by sibling species. These mosquito captures were conducted over 240 nights of surveys (Figure 1), conducted across the ILUMA Wildlife Management Area, villages immediately to the west of it, and in Nyerere National Park immediately to the east (Figure 1).

While the crude total number of adult *An. gambiae* complex mosquitoes appears to suggest higher catch on the barrier trap which is located far away from the camp (Table 1), this was because about 29% of the total collection came from only four out of 40 camps surveyed for adult mosquitoes, over only four nights of trapping: A total of 1448 adult *An. gambiae* complex mosquitoes were collected in four exceptionally large single-night samples collected at camps 14, 15, 23, and 26 (Figure 1) and supplementary file S1) in the height of the rainy season in early 2022 (Figure 2).

**Figure 2.**
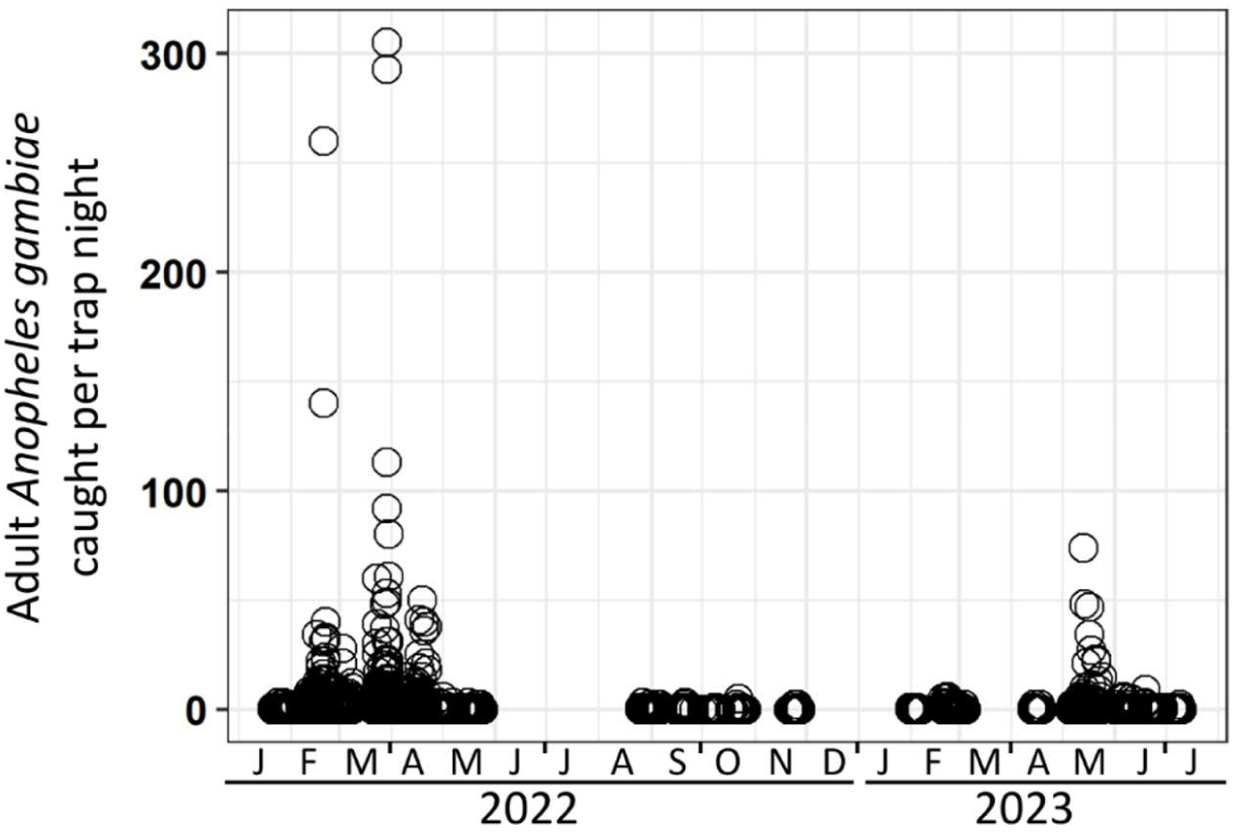
Crude number of adult *Anopheles gambiae* complex mosquitoes (Table 1) collected by light traps and an interception barrier trap in 797 trap nights over 240 nights in the surveys conducted across the ILUMA Wildlife Management Area, villages immediately to the west of it, and in Nyerere National Park immediately to the east (Figure 1).

Allowing for overdispersion (θ = 1.08 in the negative binomial GLMMs detailed in table 2, which also accounted for first order temporal autocorrelation), the mean catch of adult *Anopheles gambiae s.l.* was estimated to be much higher in the light trap placed beside the occupied tent, in which humans were constantly present throughout the night, than for any of the other light traps placed elsewhere around the camp or on the netting barrier trap placed far away from it (Table 2). No significant difference occurred between the catches obtained with any of the traps other than the light trap placed beside the occupied tent, all of which were placed further away from humans within 20 to 100 meters (p≥ 0.206). Nevertheless, for these other light trap positions, the mean catches steadily decreased as the light trap was placed further away from sleeping humans, suggesting that the attractiveness of the nearby people drew mosquitoes to these traps (Table 2), indicating all these capture methods primarily collected host-seeking mosquitoes that were attracted by humans.

**Table 2.** Estimated mean catches per trap position of adult *Anopheles gambiae* complex mosquitoes and their relative rate (RR) of capture for light traps (LT) placed in one of four distinct locations in and around the camp and one barrier trap, estimated with generalized linear mixed models as detailed in the *Methods* section. The four different positions used for LT were as follows (1) beside tent occupied by humans, (2) close to the camp, (3) in the nearby stream (4) in an open natural glade. One netting barrier interception trap was placed in an open natural glade like the fourth light trap.

| Capture method | Mean catch [95% CI] | RR [95% CI] | P value |
| --- | --- | --- | --- |
| LT Beside tent | 0.260 [0.185, 0.353] | NA | NA |
| LT In camp | 0.066 [0.042, 0.203] | 0.202 [0.138, 0.296] | <<0.0001 |
| LT Stream | 0.058 [0.036, 0.091] | 0.175 [0.119, 0.257] | <<0.0001 |
| LT Glade | 0.051 [0.032, 0.081] | 0.154 [0.105, 0.227] | <<0.0001 |
| Barrier trap | 0.053 [0.036, 0.078] | 0.159 [0.121, 0.210] | <<0.0001 |

NA: Not applicable because it is the reference group for the logistic regression model fitted.

A collection of subsamples, distributed across all camps on all occasions they were surveyed, totalling 1907 specimens of adult *An. gambaie s.l.* were tested for species identity using PCR and 76.8% (1465) were successfully amplified. Out of these amplified adult specimens, 98.7% (1446) were *An. arabiensis* and 1.3% (19) were *An. quadriannulatus*. The near complete dominance of *An. arabiensis* in these adult samples (Table 1), however, contrasted notably with the composition of the larval collections, which contained far higher proportion of *An. quadriannulatus* (Table 3). A total of 2468 larval specimens, each visually identified in the field as belonging to the *An. gambiae* complex, were collected from 248 different breeding sites, including at least one breeding site at each of the 40 camps surveyed on each survey round it was visited for (Table 3). All 2468 collected larval specimens of *An. gambiae s.l.* were subjected to molecular identification using PCR and 90.6% (2235) successfully amplified. Out of these successfully amplified specimens, 78.3% (1750) were *An. arabiensis* and 21.7% (485) were *An. quadriannulatus*.

**Table 3.** Number of breeding sites visited for larval surveys in 40 camps in the ILUMA Wildlife Management Area and neighbouring villages immediately to the west and Nyerere National Park (NNP) immediately to the east. The total number of Anopheles gambiae complex larvae collected and a breakdown of the number of each sibling species identified at each visited camp.

| Camp number and name |  | Location | Rounds<br>visited | Number of separate<br>waterbodies sampled in<br>round 4<sup>a</sup> | PCR-based species identity breakdown of<br>field-identified *Anopheles gambiae* complex larvae |  |  |  |
| --- | --- | --- | --- | --- | --- | --- | --- | --- |
|  |  |  |  |  | Total | *An. arabiensis* | *An. quadriannulatus* | Unknown<sup>b</sup> |
| 1 | Msakamba banda mbili | ILUMA | 1,2,3,4 | 1 | 7 | 7 | 0 | 0 |
| 2 | Bwawa la Msiba wa Deo | ILUMA | 1,2,3,4 | 3 | 20 | 15 | 0 | 5 |
| 3 | Bwawa la Nyati | ILUMA | 1,2,3,4 | 9 | 94 | 86 | 2 | 6 |
| 4 | Bwawa la Nandete | ILUMA | 1,2,3,4 | 9 | 102 | 87 | 1 | 14 |
| 5 | Korongo la Bundu | ILUMA | 1,2,3,4 | 9 | 70 | 62 | 1 | 7 |
| 6 | Bwawa la Namamba | ILUMA | 1,2,3,4 | 10 | 97 | 96 | 0 | 1 |
| 7 | Bwawa la Chakacheni | ILUMA | 1,2,3,4 | 10 | 104 | 76 | 0 | 28 |
| 8 | Bwawa la Njuju | ILUMA | 1,2,3,4 | 10 | 80 | 75 | 3 | 2 |
| 9 | Bwawa la Chamvi | ILUMA | 1,2,3,4 | 7 | 42 | 39 | 2 | 1 |
| 10 | Bwawa la Miembeni | ILUMA | 2,3,4 | 10 | 110 | 102 | 8 | 0 |
| 11 | Kisima cha Seba | ILUMA | 1,2,4 | 7 | 35 | 32 | 1 | 2 |
| 12 | Bwawa la Maya | ILUMA | 1,2,3,4 | 1 | 14 | 14 | 0 | 0 |
| 13 | Bwawa la Mrope | ILUMA | 1,2,4 | 2 | 9 | 6 | 0 | 3 |
| 14 | Mikeregembe | ILUMA | 1,2,3,4 | 1 | 4 | 4 | 0 | 0 |
| 15 | Mdalangwila | ILUMA | 1,2,3,4 | 4 | 14 | 14 | 0 | 0 |
| 16 | Bwawa la Mnyuamachi | Village | 1,2,3,4 | 9 | 72 | 58 | 0 | 14 |
| 17 | Tuliza Moyo | Village | 1,2,3,4 | 9 | 89 | 83 | 0 | 6 |
| 18 | Mavimba Porini | Village | 1,2,3,4 | 7 | 42 | 41 | 0 | 1 |
| 19 | Bwawa la Selesusi | Village | 1,2,3,4 | 10 | 72 | 69 | 1 | 2 |
| 20 | Makingi | Village | 1,2,3,4 | 6 | 49 | 47 | 0 | 2 |
| 21 | Bwawa la Mpunga | ILUMA | 1,2,3,4 | 3 | 9 | 9 | 0 | 0 |
| 22 | Kisaki | Village | 2,3,4 | 10 | 71 | 67 | 0 | 4 |
| 23 | Uwanja wa Ndege | Village | 2,3,4 | 10 | 64 | 63 | 0 | 1 |
| 24 | Bwawa la Mkwajuni | ILUMA | 2,3,4 | 9 | 86 | 79 | 0 | 7 |
| 25 | Bwawa la Mamba Luhogi | ILUMA | 2,3,4 | 10 | 88 | 86 | 1 | 1 |
| 26 | Funga | ILUMA | 2,3,4 | 7 | 69 | 5 | 0 | 64 |
| 27 | Bwawa la Mlenda | ILUMA | 2,3,4 | 5 | 43 | 29 | 2 | 12 |
| 28 | Bwawa la Semka | ILUMA | 2,3,4 | 3 | 14 | 10 | 3 | 1 |
| 29 | Kambi ya Simba | NNP | 3,4 | 0 | 0 | 0 | 0 | 0 |
| 30 | Bwawa la Kiboko Zanzibar | NNP | 3,4 | 3 | 4 | 0 | 3 | 1 |
| 31 | Zanzibar | NNP | 3,4 | 10 | 99 | 41 | 58 | 0 |
| 32 | Bwawa la Moto | NNP | 3,4 | 10 | 76 | 53 | 20 | 3 |
| 33 | Kambi ya Mamba | NNP | 4 | 10 | 53 | 32 | 19 | 2 |
| 34 | Kambi ya Machuma | NNP | 4 | 10 | 82 | 38 | 41 | 3 |
| 35 | Serengeti Ndogo | NNP | 4 | 10 | 120 | 50 | 67 | 3 |
| 36 | Kambi ya Makutano | NNP | 4 | 9 | 81 | 44 | 24 | 13 |
| 37 | Kambi ya Mawe | NNP | 4 | 10 | 203 | 48 | 151 | 4 |
| 38 | Shughuli kubwa | NNP | 4 | 10 | 83 | 4 | 71 | 8 |
| 39 | Bwawa la Chatu | NNP | 4 | 1 | 4 | 1 | 2 | 1 |
| 40 | Bwawa la Umeme | NNP | 4 | 10 | 93 | 78 | 4 | 11 |
| **Grand total** |  |  |  | **284** | **2468** | **1750** | **485** | **233** |

<sup>a</sup> While different breeding sites were sampled at each camp in rounds 1 to 3, the larvae collected were stored in a single tube rather than using separate tubes for each breeding site, as per round 4.
<sup>b</sup> Failed to amplify by PCR

The adult catches therefore clearly contained a higher proportion of *An. arabiensis* than larval samples (χ^2^= 311.4, d.f.= 2, p<<0.0001). Given that humans are known to be one of the two strongly preferred hosts of *An. arabiensis* (Jones, 1980; Killeen et al., 2001; White et al., 1972), and the most effective place to put light traps was clearly that closest to humans (Table 2), logistic regression analysis was applied to quantify such apparent bias towards collecting *An. arabiensis* in the adult samples and assess whether the capture method influenced sibling species composition.

No difference was observed in the composition of *An. arabiensis* collected in the light trap located farthest away from people in an open glade and the larval sample (Table 4 and Figure 3). Indeed, examining figure 3, the fitted regression model and the line of equivalence essentially overlap each other, suggesting similar proportions of *An. arabiensis* in both larval collections and adult catches with light traps placed far away from people in the open natural glade (Figure 3). On the other hand, the same logistic regression model suggests similar proportions of *An. arabiensis* in the light traps located close to the camp and in a nearby streambed, both of which are almost significantly greater (Approximately 6 and 8-fold, respectively) than that in the larvae collections (Table 4 and Figure 3). Similarly, for both the barrier trap in an open glade and the light trap placed beside a human-occupied tent, *An. gambiae s.l.* catches were almost exclusively composed of *An. arabiensis* and clearly different from the larvae collections, especially in locations where the latter was dominated by *An. quadriannulatus* (Figure 3 and Table 4).

**Figure 3:**
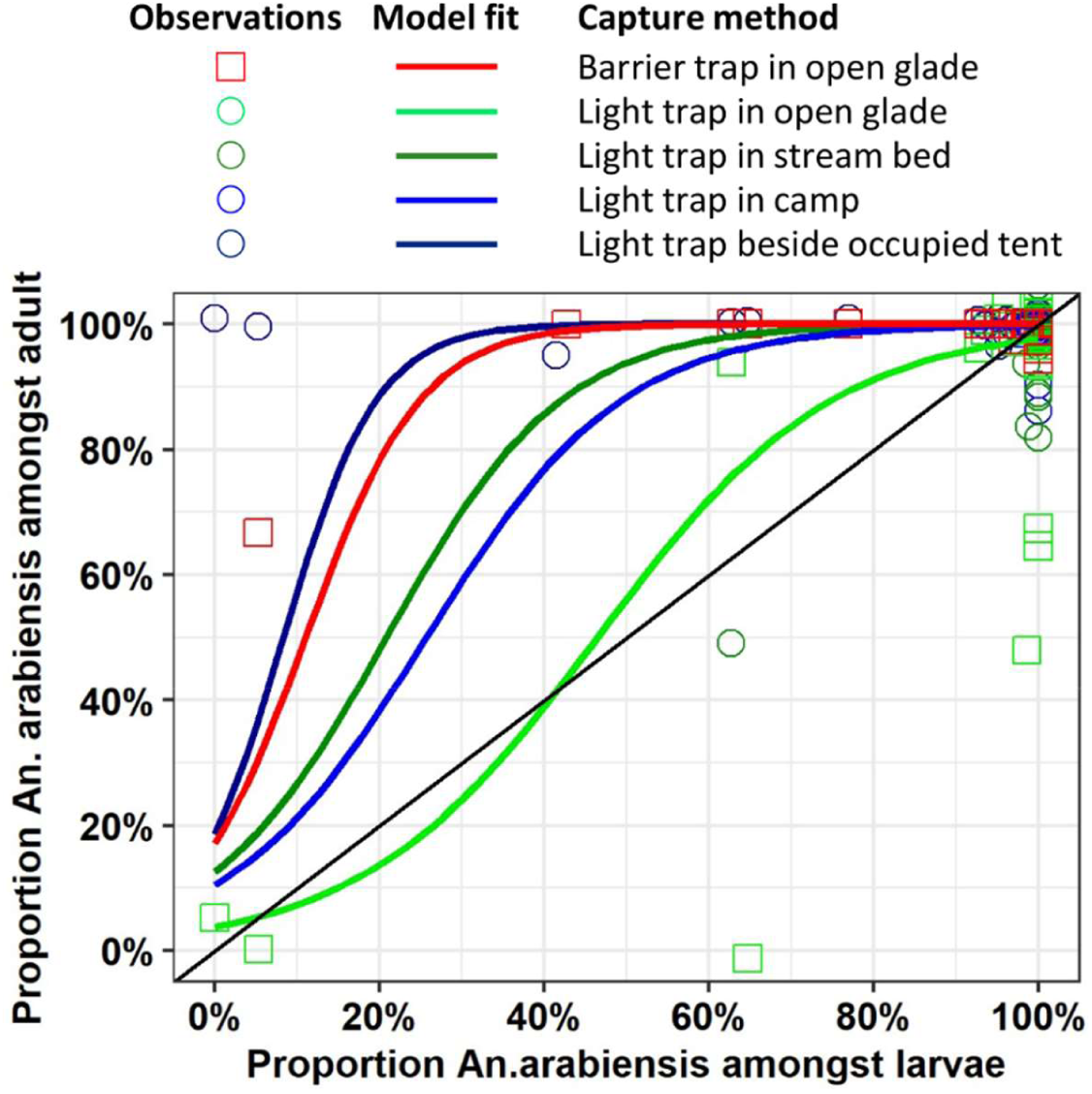
A visual summary of the observed proportion of PCR-identified *Anopheles arabiensis* in both adult and larvae samples plus fitted binomial logistic regression model curves for each capture method used during surveys of adult and larvae mosquitoes in ILUMA Wildlife Management Area, villages immediately to the west, and in Nyerere National Park immediately to the east (Figure 1).

**Table 4.** Proportions of adult and larvae samples of *Anopheles gambiae* mosquitoes that were identified as *Anopheles arabiensis*, together with the odds ratio (OR) and 95% confidence interval (CI) for being identified as this sibling species by PCR for each adult capture method compared with the larval sample, as estimated by logistic generalized linear mixed modelling. The four different positions used for placement of a light traps (LT) were (1) beside tent occupied by humans, (2) close to the camp, (3) in the nearby streambed (4) in an open natural glade, and the one netting barrier interception trap was also placed in an open natural glade.

| Capture method | Proportion *An. arabiensis* ± SE [n/N] | OR [95% CI] | P value |
| --- | --- | --- | --- |
| Larvae | 78.3 ± 0.9% [1750/2235] | NA | NA |
| Barrier trap | 99.0 ± 0.4% [770/778] | 23.9 [7.1, 80.8] | <<0.0001 |
| LT Glade | 92.8 ± 2.8% [77/83] | 1.3 [0.3, 5.5] | 0.752 |
| LT Stream | 98.8 ± 1.2% [85/86] | 8.4 [0.8, 92.9] | 0.083 |
| LT In camp | 97.6 ± 1.7% [81/83] | 5.6 [0.8, 36.5] | 0.072 |
| LT Beside tent | 99.5 ± 0.3% [433/435] | 36.2[5.6, 235.9] | 0.0002 |

NA: Not applicable because it is the reference group for the logistic regression model fitted.

In a *post hoc* GLMM analysis similar to table 4 but using barrier traps as the reference group, the odds ratio (OR) [95% confidence interval (CI)] for a specimen being *An. arabiensis* was 0.05 [0.01, 0.25] for the light trap in an open natural glade and 0.04 [0.01, 0.14] larval collections (p ≤ 0.0002 in both cases). However, no significant difference was observed between the light traps placed around the camp, in a stream bed or beside an occupied tent and the barrier trap (p≥0.109). Interestingly, when the light trap located in the open natural glade was chosen as the reference group, the odds of a specimen being *An. arabiensis* was significantly higher for the barrier trap that was similarly placed in an open area far away from the camp (OR [95% CI] 18.8 [4.0, 89.6], p = 0.002) but not for the light traps placed around the camp or in a nearby stream bed (p≥ 0.143).

Regardless of location or type of trap used, the number of *An. quadriannulatus* adults captured was consistently very low (Table 1), even inside NNP where 94.8% (460/485) of the larval collections were identified as *An. quadriannulatus*. Furthermore, the odds of an adult *An. gambiae* complex specimen caught in a light trap beside a human-occupied tent being *An. arabiensis* was 36 times higher than for the larvae collected from water bodies at the same time and place (Table 4). Therefore, this most efficient of all the trapping methods appears to be strongly influenced by the proximity of humans because it catches consistently high proportions and absolute numbers of adult *An. arabiensis* (Table 1), even when the local larval population is heavily dominated by *An. quadriannulatus* (Figure 3). A similar pattern of apparent attraction of and enrichment for *An. arabiensis* was observed for the netting barrier interception trap in an open glade (Tables 1, 2 and 4, Figure 3), suggesting that regular visits by the human research team to this device enhanced its efficiency by drawing this sibling species to the location where it was set up.

## DISCUSSION

In this study, the relative attractiveness of humans to the notorious vector of residual malaria transmission *An. arabiensis*, when compared to its non-vector sibling species *An. quadriannulatus*, was estimated based on the relative abundance of these two species in human-baited adult catches compared with samples of larvae and adults caught in non-human-baited traps placed away from people. First, however, these experiments demonstrated that human-baited light traps selectively attracted more adult *An. gambiae* complex mosquitoes than unbaited traps. Breaking these adult catches down to sibling species, the increased numbers of *An. arabiensis* caught was associated with the proximity of light traps to humans. In stark contrast, catches of wild adult *An. quadriannulatus* were consistently and remarkably sparse, regardless of survey location, trap design, or proximity to humans, even though this species dominated larval populations in well conserved, essentially undisturbed natural ecosystems. Apart from the light trap placed close to the human-occupied tent, all other light traps caught similar, much lower numbers of mosquitoes from this complex overall and *An. arabiensis* specifically.

Furthermore, light traps placed beside an occupied tent had 36 times higher odds of collecting *An. arabiensis*, rather than *An. quadriannulatus*, implying the former species is strongly attracted to humans but not the latter. Interestingly, adult catches made with the netting barrier interception trap exhibited a similar 24-fold enrichment of *An. arabiensis* relative to larval samples collected at the same time and place, presumably because this trap was actively visited by at least three people once every hour from 7 pm to midnight and then once again at 5:30 am in the morning. On each occasion, these team members spent about 10 minutes inspecting the trap and collecting mosquito resting on it, accidentally constituting a *de facto* human bait it seems. Overall, these observations consistently imply that people are strongly attractive *to An. arabiensis,* which is known to primarily rely on blood from humans and cattle (Jones, 1980; Killeen et al., 2001; White, 1974; White et al., 1972), but not to its sibling species *An. quadriannulatus*, which is generally considered a highly zoophagic species (Gillies & Coetzee, 1987; Pates et al., 2006).

Since the most recent taxonomic revision *An. quadriannulatus*, resulting in species A retaining this original name while species B was renamed as *An. amharicus* (Coetzee et al., 2013), east African populations are classified as the former and referred to as such throughout this article. Consistent with these findings that this Tanzanian population of *An. quadriannulatus* essentially ignore people as a potential blood source, complementary analyses of the same larval species composition data presented herein demonstrated a clear association of *An. quadriannulatus* with signs of activity (Duggan et al., 2024) by impala and to a lesser extent bush pig (Walsh et al., In preparation), indirectly implicating these wild ungulates as the preferred blood host of this poorly understood sibling species within the *Anopheles gambiae* complex (Walsh et al., In preparation). While some olfactometer experiments reported possible anthropophagic behaviour of this species (Pates et al., 2001), this laboratory study used laboratory colonies reared for over three years on human blood, which may well have influenced their innate behavioural preferences. In contrast, the field-based relative attraction estimates reported in table 2 for this wild *An. quadriannulatus* population in southern Tanzania indicate that it is so unresponsive to humans that it seems very unlikely that it could mediate self-sustaining levels of malaria transmission in its own right, even if it were fully physiologically competent as a host for human malaria parasites (Figure 4).

**Figure 4.**
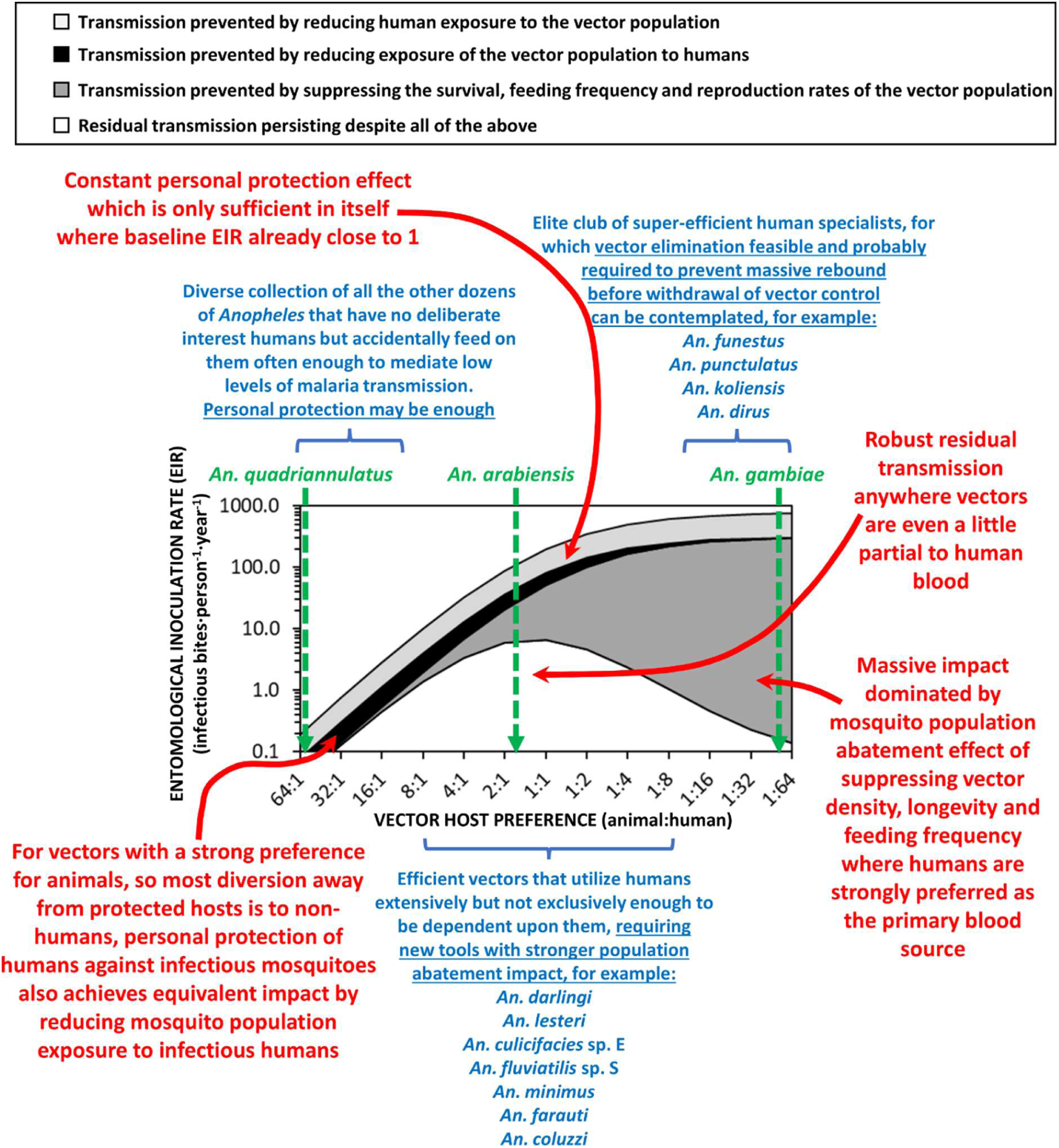
A schematic illustration of how malaria transmission intensity and responsiveness to personal protection varies according to vector preference for animals and/or humans (Killeen, 2014), and where three sibling species from the *Anopheles gambiae* complex fit into that spectrum. The relative preferences of these three *An. gambiae* complex sibling species were either estimated directly in absolute terms for *An. arabiensis* and *An. gambiae* by a previous study (Killeen et al., 2001) or extrapolated from the former estimate based on the comparative estimate for *An. quadriannulatus* versus *An. arabiensis* in relative terms that is reported herein (Table 4). The simulations were implemented as previously detailed (Killeen, 2014; Kiware et al., 2012), except that the overall impact of a personal protection measure, with equivalent deterrent and insecticidal properties to a typical modern long-lasting insecticidal net, are presented broken down by contributing underlying mechanism, and an Allee effect was incorporated (Killeen et al., 2013). Adapted from (Killeen et al., 2017).

These field observations, and the simulation analysis presented in figure 4, therefore appear consistent with the information that this species generally plays no significant role in malaria transmission (Coetzee et al., 2000; Gillies & Coetzee, 1987; Hunt et al., 1998; Lobo et al., 2015). Other than one unusual *An. quadriannulatus* population in southern Zambia (Lobo et al., 2015), which feeds on humans indoors at night (Seyoum et al., 2012) and seems remarkably vulnerable to indoor residual spraying of insecticides (Chinula et al., 2018), no wild-caught *An. quadriannulatus* has ever been incriminated as a malaria vector (Coetzee et al., 2000; Hunt et al., 1998; Lobo et al., 2015; Takken & Knols, 1999). Although one laboratory colony of *An. quadriannulatus* has exhibited some susceptibility to laboratory-cultured *Plasmodium falciparum,* it has nevertheless been found to be less physiologically competent as a vector as a colony of its sister species *An. gambiae* (Takken et al., 1999). Furthermore, some *An. quadriannulatus* populations may be outright refractory to infection with malaria parasites (Habtewold et al., 2008).

The evolutionary history of *An. arabiensis* and *An. quadriannulatus* is intricately tied to the broader evolutionary dynamics of the *An. gambiae* complex of morphologically indistinguishable sibling species, which appears to have diverged from each other on timescales that match well to those of human evolution across Africa (Fontaine et al., 2015; Neafsey et al., 2013; Neafsey et al., 2015). *An. arabiensis* appears to have diverged from *An. quadriannulatus* approximately 1.28 million years ago, and both from most of the other sibling species in the complex approximately 1.85 million years ago (Fontaine et al., 2015) through a combination of ecological pressures and geographical isolation (Bouafou et al., 2024; Schmidt et al., 2021; Soghigian et al., 2023) As a result, *An. arabiensis* has become adapted to a variety of habitats, ranging from moist savannahs through to semi-arid sahel, allowing it to become widely distributed across most of sub-Saharan Africa (Coetzee et al., 2000; Sinka et al., 2012). *An. quadriannulatus* on the other hand, is considered to be one of the more ancestral sibling species within the complex that retained several primitive traits including strong preference for non-human hosts (Ayala & Coluzzi, 2005; Thawornwattana et al., 2018).

Indeed, a complementary study of the same larval data as those presented herein indicate that the distinct innate host preferences of these two sibling species as adults, combined with fine scale variations in relative availabilities of their different preferred hosts at various location, can decisively tip the competitive balance between them one way or another (Walsh et al., In preparation). While *An. arabiensis* and *An. quadriannulatus* co-exist within essentially undisturbed natural ecosystems deep inside NNP, the latter usually dominates such locations, especially where impala are abundant (Walsh et al., In preparation). On the other hand, the apparent two-pronged specialization of *An. arabiensis* on human and cattle (Killeen et al., 2017; Killeen et al., 2001; White et al., 1972) allows it to outcompete *An. quadriannulatus* by competitive displacement (DeBach, 1966) in places wherever these hosts occur at even quite low densities (Walsh et al., In preparation).

Some useful methodological insights for informing entomological study design may also be gleaned from our findings. It makes intuitive sense that light traps placed near human-occupied tents would catch more anthropophagic mosquitoes than would otherwise be the case, and that was indeed the intended purpose of doing so from the outset. However, it was not envisaged beforehand that the netting barrier trap would yield similarly selective samples, comprised almost exclusively of *An. arabiensis* that were presumably attracted to the human members of the research team when they visited the trap to conduct collections. This relatively new design of interception trap (Burkot et al., 2013; Davidson et al., 2018; Keven et al., 2019; Tedrow et al., 2019) was chosen, and deliberately placed at a distance from the camp the team stayed at, specifically to obtain a relatively unbiased sample of mosquitoes dispersing in search of not only blood hosts, but also resting and oviposition sites. However, the almost exclusive preponderance of unfed *An. arabiensis* mosquitoes in samples obtained with this method, even in locations where larval populations were dominated by *An. quadriannulatus*, tell a different story: It seems highly likely that the research team inadvertently became *de facto* human baits every time they visited the device to check for mosquitoes. Caution may therefore be required when interpreting the results of studies using this sampling method to study mosquito population densities, species composition or bloodmeal origins, with biases towards sampling of anthropophagic mosquitoes considered as an important potential caveat. Similarly, it is also likely that experimentally controlled host preference studies with devices like mosquito electrocuting traps that are visited by the research team members on an hourly basis to collect mosquito samples (Katusi et al., 2022; Meza et al., 2019), may exaggerate catches of anthropophagic mosquitoes, even for traps that are baited with live animals. It is therefore possible that recent evidence of modest but surprizing levels of zoophagy in species like *An. funestus*, which were previously thought to be strictly anthropophagic (Finney et al., 2021; Kahamba et al., 2022), may be artefactual to at least some extent. Note also that the relatively low mean catches obtained with the netting barrier interception trap used in this study, once overdispersed spatiotemporal variation was accounted for (Table 2), despite clear enrichment for *An. arabiensis* (Table 4, Figure 3) that were presumably attracted to the collectors, seems to suggest it is probably a relatively inefficient method for catching such anthropophagic mosquitoes when they are host seeking.

## CONCLUSIONS

In conclusion, this study highlights the contrasting behaviour and host preference of *An. arabiensis* and *An. quadriannulatus.* The former demonstrates a strong attraction to humans while the latter which predominated the larval population in essentially undisturbed conserved ecosystems is non-responsive. These findings are consistent with the known behavioural ecology of both species and with those of a complementary study demonstrating that competitive relationships between these two sibling species are modulated by the relative abundances of their distinct preferred hosts around suitable larval habitats (Walsh, 2023; Walsh et al., In preparation). This study provides insights into the distinct behaviours and ecological preferences of *An. arabiensis* and *An. quadriannulatus*, which have important implications for malaria control strategies. Targeted interventions that consider species-specific behaviours, competitive dynamics, and host preference can enhance the effectiveness of malaria control programs. Additionally, the appropriate use trapping techniques can improve surveillance and provide valuable data for informing public health interventions. While light traps placed beside human-occupied tents represent a relatively efficient means of selectively capturing anthropophagic mosquitoes outdoors, unbaited light traps placed far away from people appear to give unbiased representations of larval population composition but do so with very low efficiency. Entomological survey teams visiting netting barrier devices appear to attract anthropophagic mosquitoes, practically turning them into human-baited interception traps with relatively modest capture efficiency.

## Funding Information

This study was primarily supported by an AXA Research Chair award to GFK, jointly funded by the AXA Research Fund and the College of Science, Engineering and Food Sciences at University College Cork. Supplementary funding for field equipment was kindly provided by Irish Aid through micro-project grant (Number IA-TAN/2022/144), awarded to DRM and administered by the Embassy of Ireland in Tanzania. Open access publication was funded and facilitated through the ongoing agreement between John Wiley & Sons, Inc. and the IReL consortium of Irish research libraries.

## AUTHOR CONTRIBUTIONS

Deogratius R. Kavishe: Conceptualization, investigation, methodology, formal analysis, writing original draft of the manuscript. Katrina A. Walsh, Rogath V. Msoffe, Lily M. Duggan and Lucia J. Tarimo: Implementation, review, editing, validation. Fidelma Butler: review, editing, validation. Nicodem J. Govella: Methodology, review, editing, and validation. Emmanuel Kaindoa: review, editing and validation. Gerry F. Killeen: Acquisition of fund, Conceptualization, investigation, visualization, methodology, formal analysis, review, editing and validation. All authors read and approved the final submitted version of the manuscript.

## ACKNOWLEDGEMENTS

The authors wish to thank the Village Game Scouts of ILUMA WMA, for their hard work and participation in field activities. We also thank all the governance, management, and stakeholder communities of the ILUMA WMA for all their collaboration and kind assistance over the course of the study. Furthermore, we thank Mr Frederic Masanja, Prof Honorati Masanja, Ms Catherine Ringo, Mr Fadhili Sango, Ms Elaine Kelly, Dr Ronan Hennessy, Ms Leonie O’Doherty, Ms Sonia Montero and Prof Sarah Culloty for all the essential institutional support provided by the Ifakara Health Institute and University College Cork over the course of the study. A very special word of thanks is due to our recently deceased friend and colleague, Mr Octavian Malopola, without whom this work could never have begun, much less completed safely and successfully.

## CONFLICT OF INTEREST STATEMENT

The authors declare no conflicts of interest.

## DATA AVAILABILITY STATEMENT

**Supplementary file 1;** Table S1: The number, name, location, and ecological characteristics of each camp location, together with the circuit to which it was assigned and the number of times it was surveyed https://zenodo.org/doi/10.5281/zenodo.10946752

**Supplementary File 2**; Then an experimental design form template (ED1) from a standardised mosquito collection informatics platform https://zenodo.org/records/12800114

**Supplementary File 3;** Larval survey from https://zenodo.org/records/12800143

## ORCID

Deogratius R. Kavishe https://orcid.org/0009-0007-6832-1440

Gerry F. Killeen https://orcid.org/0000-0002-8583-8739

## Notes

https://zenodo.org/doi/10.5281/zenodo.10946752

https://zenodo.org/records/12800114

https://zenodo.org/records/12800143

### Competing Interest Statement

The authors have declared no competing interest.

